# Toll-like receptor 2 stimulation augments esophageal epithelial barrier integrity

**DOI:** 10.1101/539759

**Authors:** Melanie A. Ruffner, Li Song, Kelly Maurer, Lihua Shi, Margaret C. Carroll, Joshua X. Wang, Amanda B. Muir, Jonathan M. Spergel, Kathleen E. Sullivan

**Author notes:** Correspondence: Melanie A. Ruffner MD, PhD, The Children’s Hospital of Philadelphia, Division of Allergy and Immunology, Abramson Research Center, Room 1216, 3515 Civic Center Boulevard, Philadelphia, PA 19104.

## Abstract

**Background:** A key concept of the hygiene hypothesis is that the microbiome modulates both epithelial barrier integrity as well as host immune responses. Defective expression of tight junction complex proteins alters this homeostatic process, and plays a role in atopic disorders including eosinophilic esophagitis. We tested the hypothesis that Toll-like receptor 2 (TLR2) stimulation improves esophageal barrier function in a cell-intrinsic manner by upregulation of TJ-protein expression using an *in vitro* model of human epithelium.

**Methods:** Pattern recognition receptor expression was assessed in esophageal epithelial cells from patients with EoE and non-EoE control patients. Functional consequences of TLR2 stimulation were investigated using human esophageal EPC2-hTERT cells in the three-dimensional air-liquid interface culture (ALI) model to evaluate transepithelial electrical resistance (TEER) and FITC-Dextran permeability. Characterization of TLR2-stimulated ALI cultures was performed by histology, immunohistochemistry, western blotting and chromatin immunoprecipitation.

**Results:** TLR2 stimulation increased TEER (1.28 to 1.31-fold) and decreased paracellular permeability to FITC-Dextran. Notably, TLR2 stimulation-induced increases in TEER were abolished by treatment with anti-TLR2 blocking antibody. Tight junction complex proteins claudin 1 and zonula occludens 1 were increased following TLR2 stimulation, and chromatin immunoprecipitation analysis demonstrated significant increase in histone 4 acetyl binding at the *CLDN1* enhancer and promoter following zymosan treatment, implying the occurrence of durable chromatin changes in the esophageal epithelium.

**Conclusions:** Our findings reveal that the TLR2 pathway may play a regulatory role as a mechanism that maintains epithelial barrier homeostasis in the esophagus.

## Introduction

Eosinophilic esophagitis (EoE) is an allergic disorder of the esophagus. The histology of EoE is characterized by an eosinophil-predominant infiltrate, but there is also distinctive evidence of esophageal epithelial barrier dysfunction. In EoE, the epithelial architecture shows increased basal layer hyperplasia, dilated intracellular spaces, and epithelial cell dyskeratosis (1). The consequence of this barrier dysfunction is that basal layers of the epithelium and immune cells have contact with both luminal antigens and the esophageal microbiota. Contact with food antigens stimulates secretion of Th2-type cytokines TSLP, IL-25 and IL-33 resulting in recruitment of leukocytes, including eosinophils (2–4). Additionally, EoE patients have higher rates of sensitization to esophageal microbes like *Candida* species (1,5). These studies illustrate that epithelial barrier dysfunction plays an important role in permitting antigen exposure and driving inflammation in EoE.

Tight junction (TJ) complexes regulate barrier function within the epithelium. The TJ complex is a dynamic structure comprised of transmembrane claudins, occludin, and cytosolic proteins (i.e.: zonula occludens 1, 2 and 3) which connect the TJ to the cytoskeleton (6–8). Alterations of TJ protein expression have been shown to contribute to epithelial barrier dysfunction, which is of particular consequence in eosinophilic esophagitis (1,9). Expression of occludin and claudin1 are decreased in biopsy tissue of EoE patients before and after treatment with swallowed corticosteroid (1). Further exposure to TGF-β1, which is elevated in EoE, diminished *in vitro* barrier function of immortalized esophageal epithelial cells via specific reduction in expression of the tight-junction molecule, claudin-7 (10). Additionally, the expression of ZO-1 has been shown to be significantly decreased in the lower esophagus of patients with reflux symptoms nonresponsive to proton pump inhibitor therapy who did not have evidence of EoE, illustrating that esophageal tight junctions are altered in non-EoE associated esophageal pathology (11).

In other gastrointestinal sites, innate sensing of microbial products by the epithelium has been shown to regulate epithelial barrier function. Signaling via innate immune pattern-recognition receptors on epithelial cells has been shown to modulate TJ complex protein expression, thereby modulating epithelial barrier permeability. An increasing body of evidence suggests that Toll-like receptor 2 (TLR2) signaling upregulates TJ proteins ZO-1 and claudin 1in epithelium in the airway, intestine and skin (12–17). This mechanism is thought to be one of many host defenses maintaining balance with commensal bacteria. In this study, our aim was to determine if the esophageal epithelium has mechanisms to modulate TJ complex protein expression based upon innate immune sensing via TLR2. In these studies, we utilized the three-dimensional air-liquid interface (ALI) three-dimensional epithelial *in vitro* culture model. We hypothesized that TLR2 stimulation would upregulate TJ proteins and may improve esophageal barrier function.

## Methods

### Esophageal epithelial cell culture

For this study, we used the immortalized esophageal epithelial cell line (EPC2-hTERT) (18,19). We also isolated primary esophageal epithelial cells from three pediatric control patients and three EoE patients as previously described (20). This was conducted with approval from the Children’s Hospital of Philadelphia Institutional Review Board following the US Federal Policy for Protection of Human Subjects. Patients were enrolled in the study following informed consent.

Per diagnostic criteria, all active EoE patients had eosinophil count >15 per high powered field on biopsy. Non-EoE control patients had no anomalies on esophaeal or distal histopathology, no history of immunodeficiency or other identified GI pathology. Cells were grown in keratinocyte serum-free media supplemented with bovine pituitary extract and epidermal growth factor (KSFM, Thermo-Fischer Scientific, USA). Endotoxin-free reagents and media were used for all cell culture and stimulation experiments.

### Flow Cytometry

LSR-Fortessa and FlowJo software (BD biosciences, USA) were used to examine TLR2 (11G7, Invitrogen mouse anti-human) and TLR4 (HTA125, eBiosciences/Thermofischer mouse anti-human) expression on EPC2-hTERT cells compared to isotype controls.

### Air-liquid interface (ALI) culture system

We grew confluent monolayers submerged on a 0.4 μm filter (Corning Life Sciences, USA) in keratinocyte serum-free media (KSFM, Thermo-Fischer Scientific, USA) for three days (protocol schematic, Figure 2) (21). The confluent monolayers were then switched to high-calcium concentration KSFM media (1.8 mM Ca++) for 5 days. The culture media was removed from the upper chamber to promote epithelial differentiation for days 7 to 10 (21).

### TLR2 stimulation

10 μg/ml zymosan (Infigen) and 10 μg/ml peptidoglycan from *Staphylococcus aureus* (PGN, Invivogen) were added to monolayer or basolateral chamber of ALI cultures on days 10-14. For inhibition experiments, 5 μg/ml rat anti-human polyclonal antibody to TLR2 was reconstituted per manufacturer directions and added to basolateral chambers containing 10 μg/ml zymosan (Invivogen, PAb-hTLR2).

### Transepithlial electrical resistance (TEER

We measured ALI culture TEER with a Millicell ERS-2 Voltohmmeter (Merck Millipore, USA). The final unit area resistance (Ω*cm^2^) was calculated by multiplying the measured sample resistance by the membrane area (0.33 cm^2^ for 24-well Millicell inserts). Only epithelial monolayers with TEER >200 ohms·cm^2^ on day 10 of ALI culture were used for stimulation experiments.

### Transepithelial Flux

We determined the flux of 70 kDa fluorescein isothiocyanate-dextran (FITC-dextran; Sigma, USA) from day 14 ALI cultures by adding 3 mg/mL 70kDa FITC-dextran solution to the upper culture chamber. The initial 3mg/mL upper solution was sampled to generate the standard dilution curve. The fluorescein levels in basolateral samples were detected after 4 hours on a Spectramax M5 plate reader (Molecular Devices, San Jose USA) and normalized to the mean of the unstimulated control samples.

### Histology and Immunohistochemistry

We harvested ALI on day 14. Samples were fixed in formalin, paraffin-embedded and serially sectioned. Slides were deparaffinized in xylene and rehydrated thru a series of ethanol washes. Slides were stained in hematoxylin, rinsed, then stained in eosin (Azer Scientific, USA) on a Shandon Gemini automated stainer (ThermoFisher, USA). Slides were incubated with E1 (Leica Biosystems) immunohistochemistry antigen retrieval solution for 20min. Claudin 1 (1:50; LS Bio LS-C415827; 1hour primary incubation) and ZO-1 (1:50; Sigma HPA001636; 1hour primary incubation) antibodies were used for staining with Bond Refine polymer staining kit on the Bond Max automated staining system (Leica Biosystems, Germany). All stained slides were dehydrated in ascending ethanol and xylene washes before coverslipping with cytoseal (Fisher Scientific, USA). Stained slides were digitally scanned at 20x magnification on an Aperio CS-O slide scanner (Leica Biosystems). ALI membrane thickness, number of basolateral nuclei, percent dilated intracellular space and percent strong 3,3′-Diaminobenzidine (DAB) staining intensity were calculated in Aperio imaging software.

### Western Blot

Day 14 ALI were washed in PBS and lysed in RPIA buffer containing protease cocktail. Sample protein content was normalized via Bradford assay prior to electrophoresis on a 4-12% Bis-tris gel, followed by transfer to nitrocellulose membranes. Mouse anti-human β-actin (1:1000, cell-signaling technologies 3700S), rabbit anti-human Claudin 1 (1:500, LSBio LS-C415827), and rabbit anti-human ZO-1 (1:500, HPA001636 Sigma) primary antibodies were used to probe the membrane. Secondary HRP conjugated anti-rabbit and anti-mouse antibodies (Cell signaling technologies) were used at a concentration of 1:2000. NIH ImageJ was used for semi-quantitative densitometry analysis.

### Quantitative real-time PCR (qRT-PCR)

We harvested RNA using Directzol (Zymo Research, Irvine, CA). DNA was removed by column. We used 0.2 μg of total RNA for reverse-transcription reactions using Advantage RT-for-PCR kit (TakaRa Bio USA). cDNA was used for RT-PCR. Taqman gene expression assays (Applied Biosystems, human *TLR1-10, CLEC7A, CD14),* were normalized to the *GAPDH* internal control signal and custom SYBR green primer assays (listed below) were normalized to *β-actin* control. Three independent replicates were averaged for the results presented.

### Chromatin Immunoprecipitation (ChIP)

Five million cells per condition were used to perform ChIP experiments following the protocol from Upstate Biotechnology (Lake Placid, NY) with some modifications (22). EPC2-hTERT were harvested from monolayer after 30 minutes of stimulation, treated with 1% formaldehyde for 10 min at room temperature, sonicated and immunoprecipitated overnight at 4 °C. 5 μg of the following antibodies were used: H4ac (Histone 4 pan-acetyl, Merck Millipore #06-866) anti-H3K27ac (Abcam ab4729), trimethylated H3 lysine 4 (Active Motif 39159), and anti-glutathione-S-transferase (GST) antibody (ThermoFisher 71-7500). Antibody-bound complexes were collected, eluted and extracted as previously described, (22) then RNase treated, and quantitated before analysis by qRT-PCR. The primers for the ChIP assays are listed below.

#### Primers for SYBR qRT-PCR

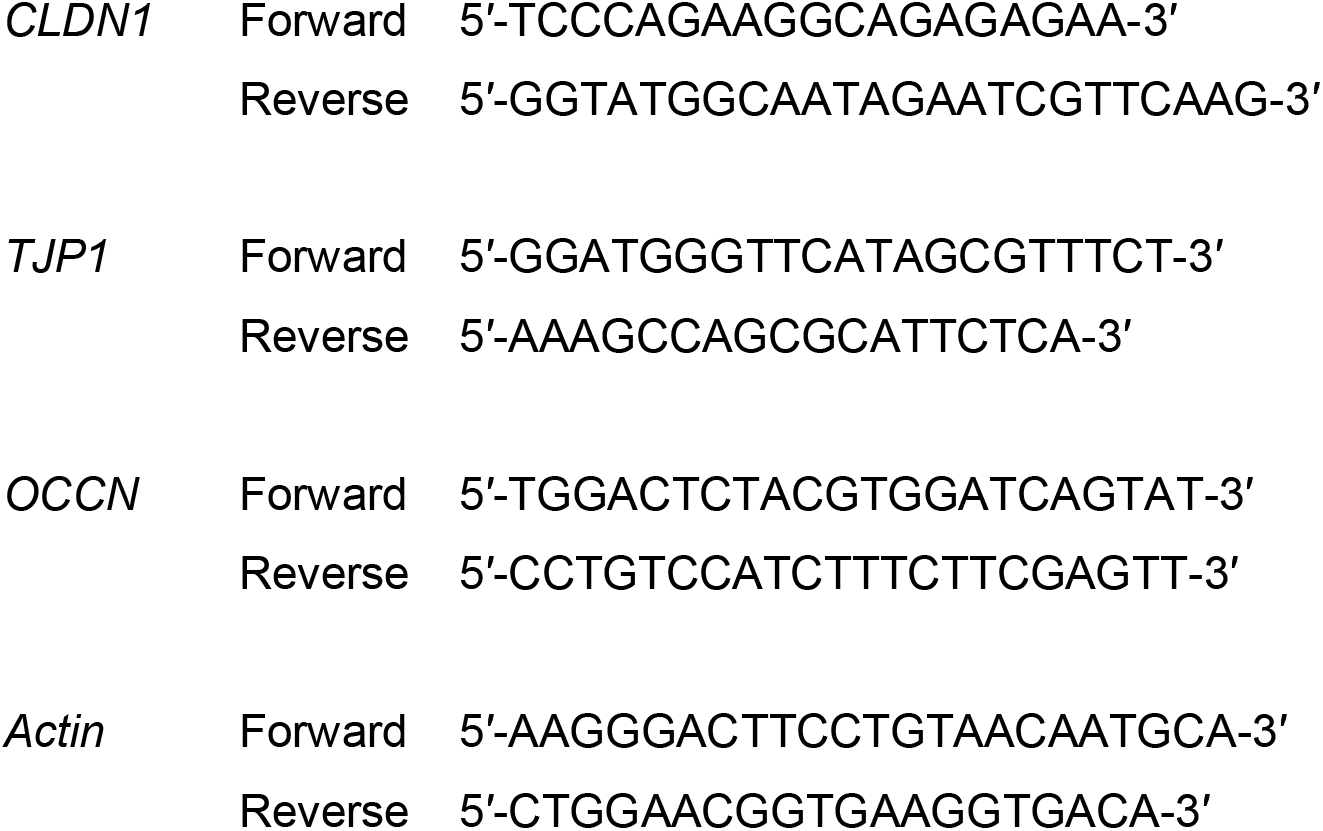

#### Primers and probes for ChIP assay

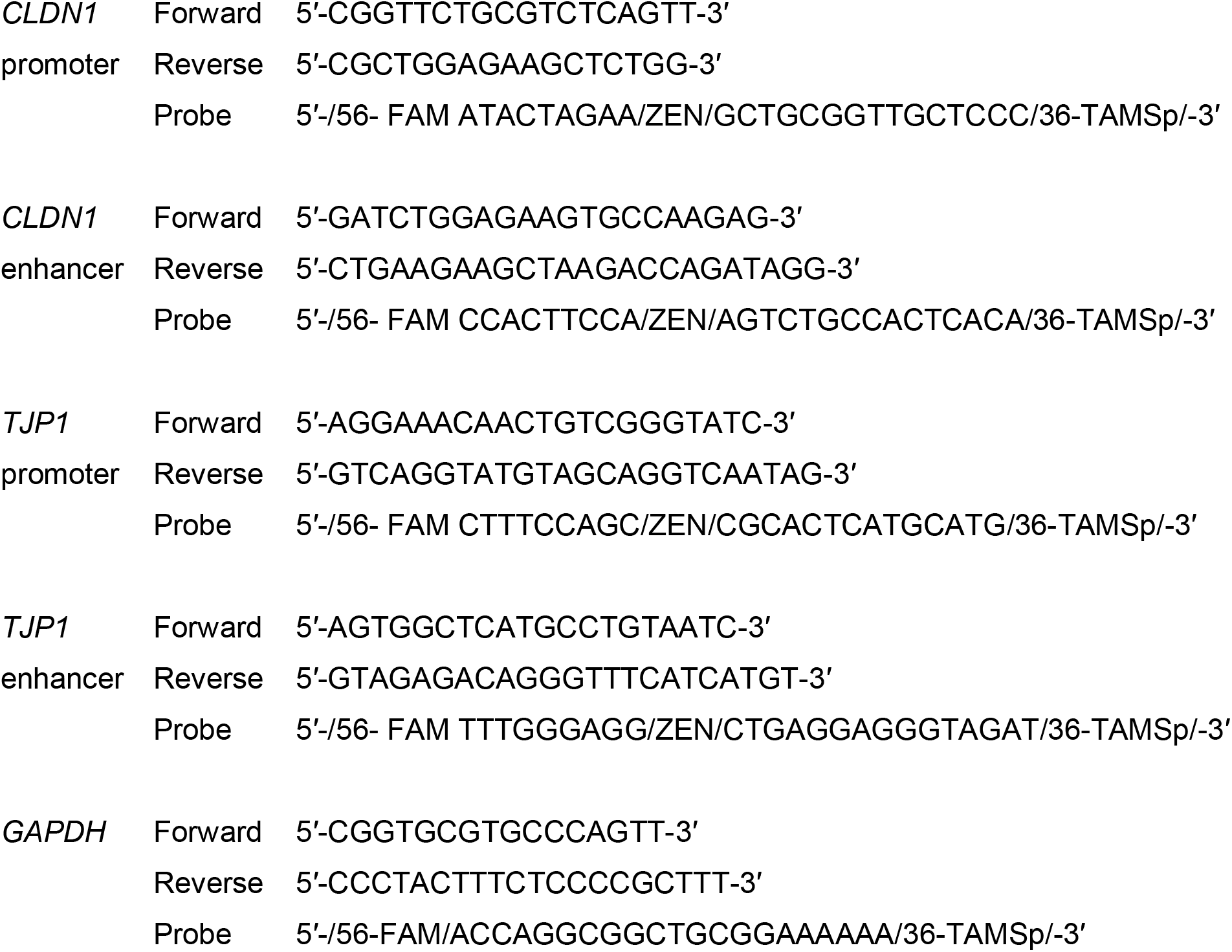

### Statistical analysis

Statistical tests were performed using GraphPad Prism version 7.0 (San Diego, CA, USA). Data are presented as mean ± SD. p < 0.05 was considered to be statistically significant.

## Results

### TLR2 and TLR3 were highly expressed in esophageal epithelial cells

We hypothesized that TLRs were involved in regulating epithelial barrier function. To test this hypothesis, we examined expression of PRRs in esophageal epithelial cells from control subjects, subjects with EoE and in the EPC2-hTERT esophageal epithelial cell line (Figure 1A). We examined gene expression patterns of TLRs 1–10, CD14 and Dectin 1 (*CLEC7A*) using qRT-PCR. Both primary esophageal epithelial cells and the EPC2-hTERT cell line express *TLR1* through *TLR6*, with minimal expression of *TLR7* through *TLR10, CD14* and *CLEC7A*. TLR2 forms heterodimers with TLR6 and TLR1, both of which were transcribed in the esophageal cells. Both primary and immortalized cell lines expressed TLR2 and TLR3 at the highest levels compared to other PRRs tested. TLR2 protein is expressed on the cell surface, and was detected on epithelial cells using flow cytometry (Figure 1B & D). In comparison, the levels of Dectin 1 staining did not exceed isotype control staining (Figure 1C). These studies demonstrate that esophageal epithelium expresses a complement of PRR and confirm cell surface expression of TLR2.

**Figure 1:**
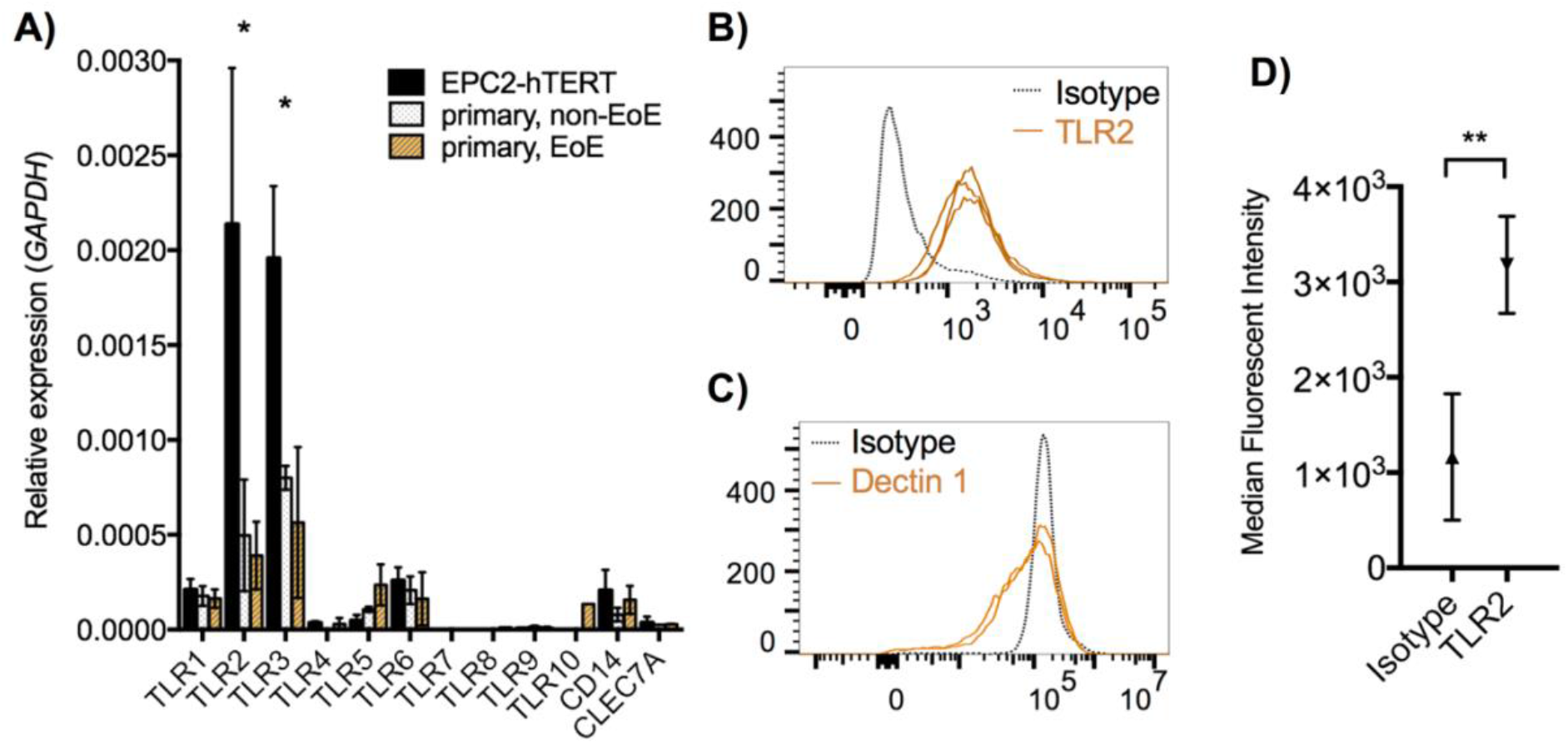
Esophageal epithelial cells express high levels of TLR2 and 3. **A)** *TLR1* thru *TLR10, CD14* and *CLEC7A* (Dectin 1) gene expression was determined by qRT-PCR in primary esophageal epithelial cell lines from patients with and without eosinophilic esophagitis (EoE) and from immortalized esophageal epithelial cell line EPC2-hTERT (* p<0.05 by ANOVA with post-hoc t-test; EPC2-hTERT vs. EoE and EPC2hTERT vs. non-EOE, mean ±SD, *n*=3 biologic replicates). **B)** EPC2-hTERT cells stained with anti-TLR2 or **C)** anti-Dectin 1 antibodies compared to isotype control antibody (black dashed line) using flow cytometry. Orange lines denote individual stained samples. **C)** Median fluorescence intensity of TLR2 versus isotype expression is shown (**p<0.001 by Mann-Whitney test, *n*=3).

### TLR2 stimulation improved epithelial barrier function in air liquid interface culture

We next set out to test the effect of TLR2 stimulation on esophageal epithelial cell barrier function. We used the three-dimensional air liquid interface (ALI) model and added TLR2 ligand or cytokine to the basolateral chamber media on days 10-14 of culture (Figure 2, schematic). Stimulation with TLR2/6 agonist zymosan (100 μg/mL) resulted in increased transepithelial electrical resistance (TEER, 28% increase, Figure 2B & D, ANOVA, Fisher LSD posthoc p<0.05 on day 14). We observed a similar magnitude of TEER increase in ALI cultures treated with peptidoglycan (PGN), which stimulates the TLR2/1 complex (figure 2B, 31% increase, p<0.001 on day 14). Using the FITC-dextran permeability assay, we observe that the basolateral concentration of FITC-dextran in zymosan and peptidoglycan are significantly decreased compared to untreated control samples at 4 hours (figure 2C). indicating improved barrier function in the TLR2-agonist treated samples.

**Figure 2:**
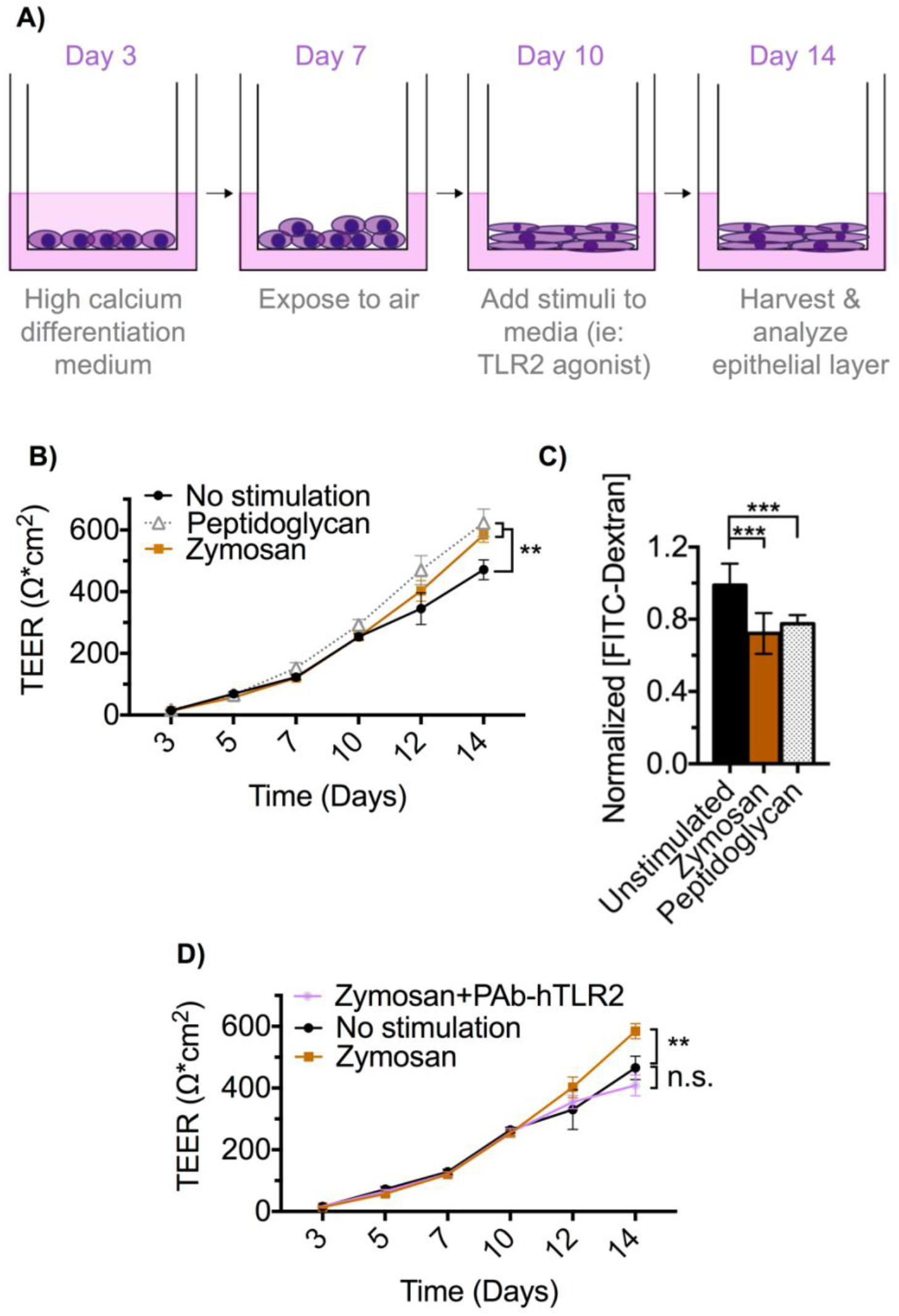
TLR2 stimulation of EPC2-hTERT in ALI culture results in improved barrier function. **A)** Schematic of ALI epithelial culture model. Culture days 1-7 allow for proliferation and initial differentiation while days 7-14 represent a terminal differentiation phase. Stimuli applied in the media of the basolateral chamber during days 10-14 can be assessed for effect on epithelial barrier terminal differentiation and function. **B)** ALI cultures were stimulated with zymosan (100 ng/mL) or peptidoglycan (100 ng/mL) from days 10-14. Zymosan and peptidoglycan treated cultures demonstrate a significant increase in TEER on day 14 when compared to unstimulated ALI cultures. Transepithelial electrical resistance (TEER) was measured by ohmmeter and final unit area resistance (Ω*cm^2^) was calculated by multiplying the sample resistance by the area of the membrane. n=10 wells per group, representative of three experiments. *** denotes p<0.001 using ANOVA followed by Fisher’s LSD test. **C)** On day 14, translocation of 70 kDa FITC-Dextran across ALI membranes into the basolateral culture chamber was decreased in the zymosan and peptidoglycan treated groups (*n*=6 wells per group, mean ±SD. Representative of two experiments. **p<0.01, and ***p<0.001 ANOVA with posthoc Fisher’s LSD). **D)** Addition of pAB-hTLR2, anti-TLR2 blocking antibody to ALI cultures stimulated with 100ng/mL of zymosan block zymosan-induced TEER increase. **p<0.01, n.s. denotes p>0.05, ANOVA with posthoc Fisher’s LSD)

### Improved barrier function following zymosan treatment is TLR2-specific

The full complement of PRR in esophageal epithelium has not been defined and their function are poorly understood. TLR2 coordinates with TLR6 to recognize zymosan, however zymosan can also be recognized by Dectin-1 as well as other potential epithelial-specific receptors such as Ephrin A2 (23). Although the EPC2-hTERT have little Dectin 1 expression, we set out to determine if the observed improvement in epithelial barrier function was TLR2-specific. To do so, we co-incubated ALI cultures with PAb-hTLR2 and zymosan (Figure 2D). PAb-hTLR2 is a polyclonal rat anti-human blocking antibody specific for TLR2 (24,25). We observed that the addition of PAb-hTLR2 into the culture media abrogated the increase in epithelial TEER seen following stimulation with zymosan, demonstrating a specific requirement for TLR2 to improve esophageal epithelial barrier function *in vitro*.

### TLR2 stimulation alters epithelial membrane architecture

We next examined epithelial architectural changes associated with zymosan and peptidoglycan treatment by histological analysis of ALI cultures (Figure 3). We observe the expected stratification of the epithelial membrane in unstimulated samples, with thinning of the epithelial membrane following TLR2 agonist treatment (24.8±5.6 μm untreated vs 17.6±6.5 μm zymosan-treated, p<0.001, and 18.7±3.9 peptidoglycan-treated p<0.05). The density of basolateral nuclei was decreased in TLR2 agonist treated cultures (Figure 3C). Dilated intracellular spaces are a characteristic finding of epithelial dysfunction in EoE, however, there were no differences amount of dilated intracellular space between treated and untreated cultures (Figure 3D). Overall, this corresponds with data obtained in our barrier function assays. Although TLR2 stimulation induces morphologic changes within the epithelium there is no indication of epithelial dilated intracellular spaces which have been demonstrated to correlate with poor barrier function (26,27).

**Figure 3:**
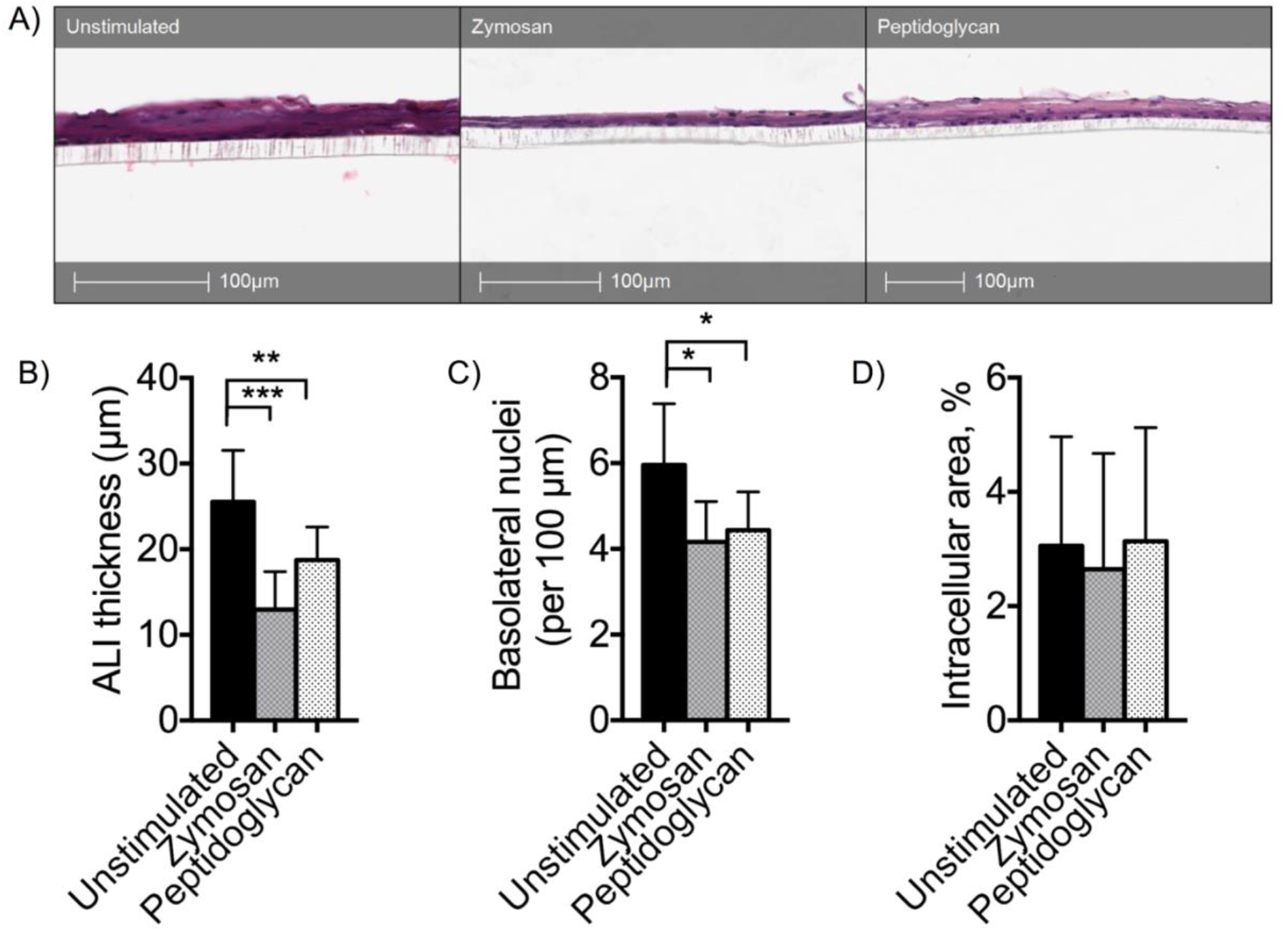
Zymosan stimulation of esophageal ALI alters epithelial morphology. **A)** ALI were harvested on day 14 of culture, fixed in paraffin then stained with hematoxylin and eosin. Images were captured at 20x. We quantified **B)** ALI thickness, **C)** basolateral nuclei (expressed per every 100 μm of basolateral membrane adjacent to the Transwell membrane) and **D)** the percent of total area dilated intracellular space quantified using Aperio software imaging algorithms (n=5 slides per condition; mean ± SD, * p<0.05, **p<0.01, *** p<0.001, ANOVA, post-hoc t-test).

### TLR2 stimulation upregulated TJ complex molecule gene expression

TJ complex proteins are the primary mediators of epithelial barrier integrity. Stimulation of TLR2 in intestinal and airway epithelium increases expression of TJ complex proteins (13,15,17). TJ proteins including claudin 1 and ZO-1 are dysregulated in EoE. In light of the finding of improved epithelial barrier function following stimulation with TLR2 agonists, we sought to better understand how TJ complex protein expression changed following TLR2 stimulation in this model. Using qRT-PCR, we assessed expression of TJ complex genes *CLDN1* and *TJPN1* (which encode claudin 1 and ZO-1, respectively) in EPC2-hTERT cells stimulated in monolayer with zymosan or peptidoglycan (Figure 4). These were significantly upregulated over the course of 24 hours.

**Figure 4:**
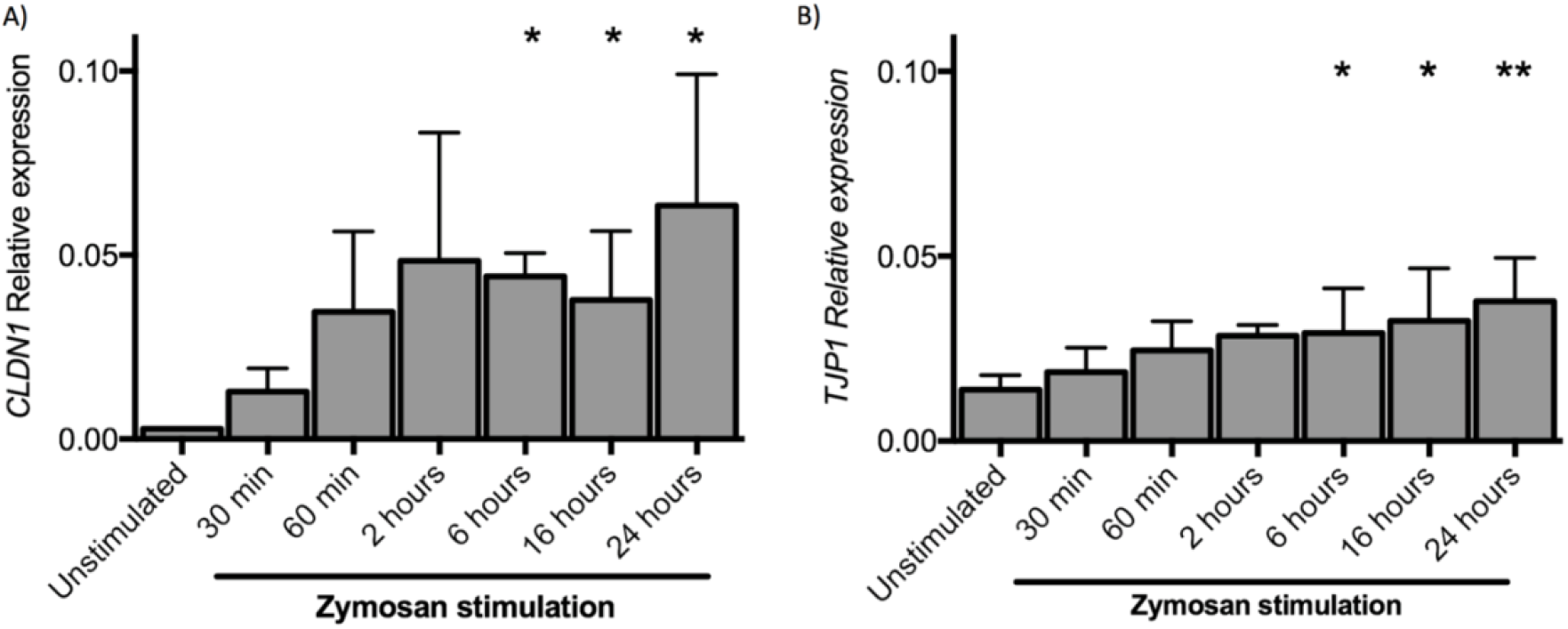
Zymosan treatment of EPC2-hTERT cells upregulates expression of cell adhesion molecules. EPC2-hTERT cells stimulated with Zymosan (100 ng/ml) in culture results in significant upregulation of **A)** *CLDN1*, and **B)** *TJP1* by 6 hours of stimulation. Expression of target mRNA is expressed relative to *GAPDH* mRNA (*n*=4; mean ± SD, * denotes p<0.05 by ANOVA with post-hoc Fisher’s LSD test)

To test the effect of TLR2 stimulation on TJ complex proteins expression in esophageal epithelial cells, we examined claudin 1 and ZO-1 protein by immunohistochemistry in ALI cultures harvested on day 14 following stimulation with zymosan or peptidoglycan (Figure 5A). Using image analysis to quantitate 3,3′-diaminobenzidine staining intensity, we confirmed that the ZO-1 and claudin 1 staining was significantly higher in zymosan- and peptidoglycan-treated ALI cultures (Figure 5B and 5C). We verified increase protein expression of claudin 1 and ZO-1 in ALI following TLR2 stimulation using western blot (Figure 5D). From these studies, we conclude that TLR2 stimulation drives increased expression of TJ proteins.

**Figure 5:**
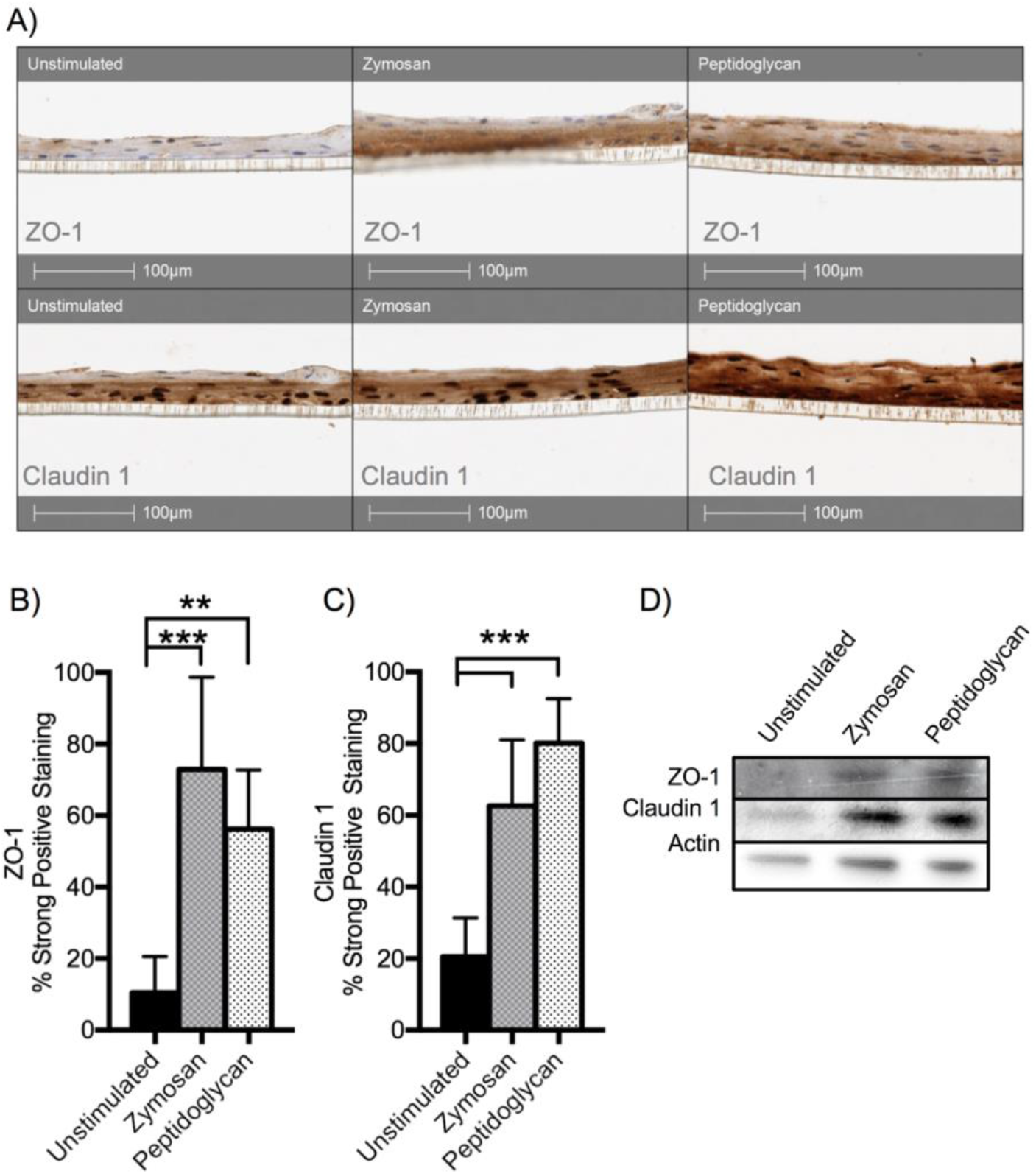
ALI cultures treated with zymosan upregulate epithelial cell-cell adhesion molecules. **A)** Immunohistochemistry of EPC2-hTERT grown in ALI culture and stimulated with zymosan and peptidoglycan. Staining for claudin 1 is diffusely upregulated but ZO-1 staining is denser in the basal layer. Using DAB staining quantification algorithm in Aperio, we find that **B)** Claudin 1 and **C)** ZO-1 staining are significantly upregulated (mean ± SD ** p<0.01 or *** p<0.001 ANOVA, posthoc LSD). **D)** Western blotting of day 14 ALI cultures. Claudin 1 and ZO-1 are upregulated in zymosan- or peptidoglycan treated samples. Results shown are representative of three experiments.

### Histone modifications in zymosan-treated epithelial cells

Modification to the chromatin regulatory landscape alters gene expression in response to changes in the cellular environment. We hypothesized TLR2 stimulation may mediate gene expression changes in tight junction complex proteins within esophageal epithelial cells. These chromatin modifications could be perpetuated across multiple divisions throughout generation of the stratified barrier. We performed chromatin immunoprecipitation (ChIP) assays of EPC2-hTERT cells following stimulation with zymosan (Figure 6) to assess for changes in the chromatin environment at the promoter and enhancer of *CLDN1* and *TJPN1*. Zymosan stimulation increased the acetylation of histone H3K27 and trimethylation of H3K4, though neither reached statistical significance (Figure 6). Acetylation of histone H4 was increased at the *CLDN1* enhancer and promoter of zymosan-treated samples. This suggests that Claudin 1 expression was regulated epigenetically by acetylation subsequent to TLR2 stimulation with zymosan.

**Figure 6:**
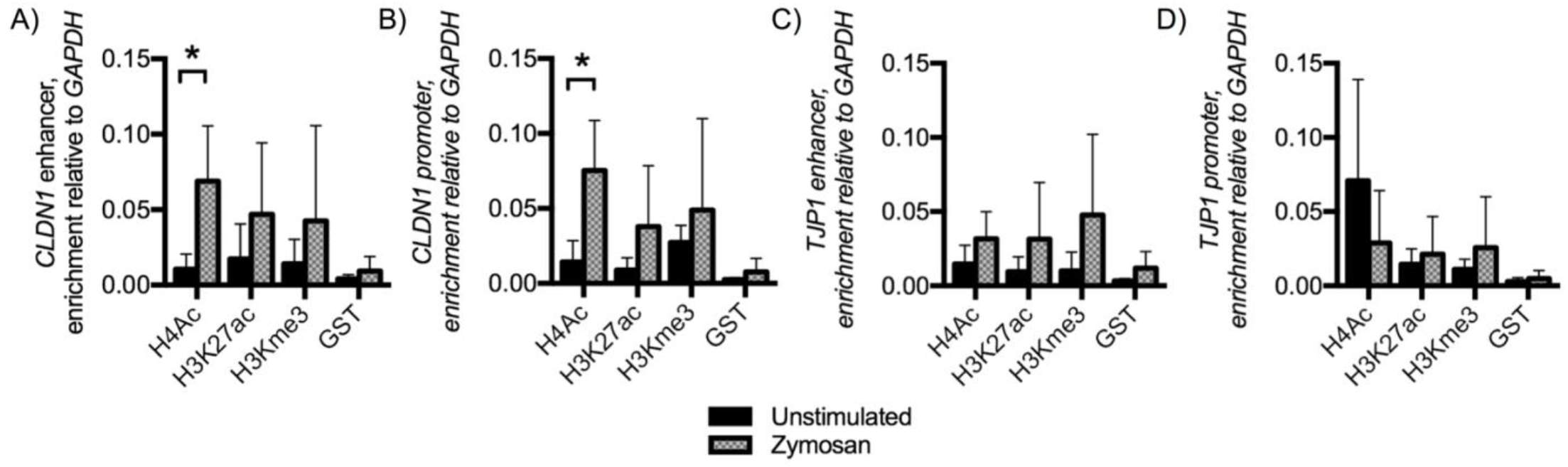
EPC2-hTERT cells stimulated with Zymosan alter chromatin at *CLDN1* and *TJP1* promoters and enhancers. After 30 min of incubation with zymosan, chromatin immunoprecipitation assays were performed using antibodies specific to H4ac, H3K27ac, H3Kme3, or control antibody, GST. PCR was performed to amplify immunoprecipitated DNA for the **A)** *CLDN1* enhancer locus (*n*=4, mean ± SD *p<0.05, t-test), **B)** *CLDN1* promoter locus, (*n*=4, mean ± SD p<0.05, t-test) **C)** *TJP1* enhancer locus (ns, n=4 mean ± SD), or **D)** *TJP1* promoter locus (ns, n=4 mean ± SD).

## Discussion

Our data suggest that recognition of lipoprotein by TLR2 is a specific mechanism that increases epithelial barrier integrity within the esophageal epithelium (Figure 2). Utilizing the air-liquid interface model with the esophageal EPC2-hTERT epithelial line, we demonstrate that TLR2 stimulation results in an increase in the expression of TJ-associated proteins claudin 1 and ZO-1 (Figures 4 and 5). EPC2-hTERT cell lines have been previously characterized, and share the same morphological characteristics as esophageal basal keratinocytes including key differentiation markers like cytokeratins and the ability to terminally differentiate *in vitro* in high extracellular calcium media or if cultured to post-confluent state (18,19,28). The esophagus is not a sterile environment, and changes in microbiome composition have been reported in patients with reflux, Barrett’s esophagus as well as EoE patients with active inflammation (29–31). Further, tissue damage from inflammation or reflux causes release of endogenous damage-associated molecular patterns from necrotic cells which can also interact with esophageal PRR (32,33). Therefore, it is plausible that changes in the milieu of PRR-ligands affecting the esophageal epithelium can occur without clinical infection. Our findings describe a mechanism for esophageal barrier modulation in response to these changes.

TJ complex molecules claudin 1 and ZO-1 are upregulated within 24 hours following TLR2 stimulation (Figure 4). Our results are consistent with findings from other epithelial types, including intestinal, airway and skin keratinocytes, where TLR2 agonism has been associated with increased expression of TJ complex molecules claudin 1 and ZO-1 (12–15,17,34). In this study we focused on the response of claudin 1 and ZO-1 because of their critical role in tight junction complex formation. We hypothesize that TLR2 stimulation induces additional alterations in esophageal epithelial cell gene expression that were not captured in our experiments. Characterizing these additional functional effects will clarify the full impact of TLR2 stimulation on the esophageal mucosa.

Extrapolating upon these findings, we hypothesize that signaling thru innate immune receptors may play a critical role in epithelial cell fate. We find that stimulation of TLR2 with zymosan results in significant enrichment of H4ac at both the *CLDN1* promoter and enhancer (Figure 6). H4ac is an activation mark with a widespread distribution at regulatory regions. TLR2-induced chromatin remodeling could be secondary to an overall shift in the balance of histone acetyltransferases and deacetylate enzymes or more likely due to targeting of chromatin modification complexes to the regulatory regions (35). Novel therapies with HDAC inhibitors could be investigated to target this pathway within the mucosal for disorders like EoE.

TLRs, including TLR2 and TLR1 are upregulated in the mucosa of patients with EoE and downregulated following treatment with six-food elimination diet (36). TLR2 single nucleotide polymorphisms (SNP) have been associated with asthma in Puerto Rican, Norwegian, Danish and Caucasian African populations (37–43), as well as development of severe atopic dermatitis in Italian and German populations (44,45). The mechanism of each SNP is not fully elucidated, however several of the SNPs decrease TLR2 signaling activity (46). Recent data confirm that EoE is a part of the atopic march, wherein the cumulative risk of EoE increases with each subsequent allergic comorbidity (47). Further studies elucidating how PRRs such as TLR2 contribute to host barrier responses may help clarify mechanisms of atopic disease including EoE.

In conclusion, we have shown that esophageal epithelial cells express low levels of *TLR 1* through *TLR6*, with highest levels of *TLR2* and *TLR3* expression. Stimulation of TLR2 with lipoproteins zymosan and peptidoglycan result in upregulation of Claudin 1 and ZO-1, resulting in enhanced epithelial barrier function. This effect is TLR2-specific and abrogated by TLR2 blocking antibody. Our data highlight a potential homeostatic role of esophageal epithelial TLR2 activity in upregulating tight junction complex protein expression. This has implications for esophageal mucosal responses to infection, colonizing microbiota and endogenous DAMPs released in the context of reflux and inflammation. Based on the apparent protective effect of TLR2 agonism on the mucosal barrier in esophageal epithelium, we conclude that epithelial innate signaling pathways may provide novel therapeutic targets for the future treatment of esophageal disorders such as EoE.

## Acknowledgements

EPC2-hTERT were the kind gift of Dr. Hiro Nakagawa. We wish to thank Drs. De’broski Herbert, Rob Rubenstein and Yann Bikard for advice regarding the ALI model, and Swathi Raman for technical assistance, and the Children’s Hospital of Philadelphia Flow Cytometry and Pathology Core Facilities.

## Author contributions

MAR conceived of the presented idea. MAR, ABM, JMS KES, developed idea, reviewed & interpreted data. ABM performed endoscopies and isolated epithelial cells. MAR, LS, LHS, MCC, KM, JXW performed experiments, analyzed & data. MAR wrote article with review and comments from all other authors.

## Abbreviations

TLR: toll like receptor
PRR-: pattern recognition receptor
ALI-: air liquid interface
EPC2-hTERT-: Primary human esophageal keratinocytes, EPC2 human telomerase immortalized
*CLDN1*: claudin 1
ZO-1: zonula occludens 1
*CAPN14*: calpain 14
TEER: transepithelial electrical resistance
KSFM: keratinocyte serum free media
PGN: peptidoglycan
GST: : glutathione-S-transferase
USA: United States of America
HAT: : histone acetyl transferase
HDAC: : histone deacetylases
SNP: : single nucleotide polymorphism

